# Sensitive detection of copy number alterations in low-pass liquid biopsy sequencing data

**DOI:** 10.1101/2025.09.22.677749

**Authors:** Lotta Eriksson, Eszter Lakatos

## Abstract

Liquid biopsies, coupled with analysis of copy number alterations (CNAs), have emerged as a promising tool for non-invasive monitoring of cancer progression and tumor composition. However, methods utilizing CNA data from liquid biopsies are limited by the low signal in the samples, caused by a low percentage of cancer DNA in the blood, and inherent noise introduced in the sequencing. To address this challenge, we developed BayesCNA, a method designed to improve signal extraction from low-pass liquid biopsy sequencing data, by utilizing a Bayesian changepoint detection algorithm. We use information of the posterior changepoint probabilities to identify likely changepoints, where a changepoint indicates a shift in the copy number state. The signal is then reconstructed using the identified partition. We show the effectiveness of the method on synthetically generated datasets and compare the method with state-of-the-art bioinformatics tools under noisy conditions. Our results show that this novel approach increases sensitivity in detecting CNAs, particularly in low-quality cases.

## 1. Introduction

Routine sampling and sequencing of patients’ tumors are crucial for enabling precision medicine. However, traditional tissue-based biopsies are invasive and repeatedly collecting representative samples is often impractical, especially for patients with metastatic cancer. Acquiring tissue samples during treatment to monitor tumor response and relapse is particularly challenging. To address the challenges with traditional biopsies, recent oncology research has shifted towards the analysis of bodily fluids for tumor-derived components, referred to as liquid biopsies [17]. Liquid biopsies are promising for longitudinal and minimally invasive monitoring of cancer dynamics, mainly by utilizing cell-free DNA (cfDNA) present in blood samples that is released by necrosis or apoptosis of the cell [15]. Tumor-derived cfDNA is correlated with the stage of the disease [6, 8], offers the same diagnostic potential as tissue-based biopsies [14], while being less invasive, and produces a comprehensive overview of systemic disease.

Copy number alterations (CNAs) are structural variations that result in discrete gains or losses of large genomic regions. CNAs are prevalent in human cancers and play an important role in cancer progression by activating oncogenes and inactivating tumor suppressors [18]. CNAs are often exclusive to tumor cells, making them a good tool for cancer detection and tracking. CNAs can be detected using cheap lowpass whole genome sequencing (lpWGS), typically performed at coverages of 0.001-0.1x. Hence, liquid biopsies coupled with the analysis of CNAs are a promising tool for non-invasive monitoring of cancer progression and tumor composition. However, tumor cells are not the only contributor to the DNA pool in the blood, where healthy DNA often comes in a high proportion. Since healthy DNA does not carry the genomic alterations present in cancer cells, the signal in the sample is reduced, making detection of CNAs challenging. Although a wide range of methods are available for CNA detection in tissue biopsies [11, 7, 10, 12, 9], including lpWGS samples, these are hampered by the low signal commonly observed in liquid biopsies and tend to break down in low coverage and/or low purity scenarios. To address these shortcomings, we developed BayesCNA, a method that utilizes Bayesian changepoint detection (BCP) for sensitive detection of CNAs in low-quality liquid biopsy sequencing data.

Changepoint detection is the problem of identifying times or spatial positions where the probability distribution of a stochastic process or time series changes abruptly. Changepoint detection is used in various fields, including medicine and finance; see, e.g. [5, 4]. In the setting of CNA detection, changepoints correspond to the boundaries of segments, contiguous regions of the genome with identical copy number state. Accurate detection of changepoints is therefore crucial for correct partitioning of the genome and identification of altered copy number regions. Information about the location of CNAs is important in longitudinal study of cancer dynamics, where we are interested in studying how segments change over time, for example, to estimate the subclonal composition of a tumor at different timepoints [16]. Changepoint detection has been widely used in the analysis of copy number alterations, and the most popular approaches are circular binary segmentation (CBS) [2] and models based on a Hidden Markov Model (HMM) [12]. However, CBS struggles to detect changepoints when the change in mean is small, and when the changed segment is short. HMM-based methods, on the other hand, require a predefined set of possible states, which are often unknown a priori. An alternative to traditional changepoint methods is Bayesian changepoint detection (BCP), introduced by Barry and Hartigan [1] as a Bayesian approach to the changepoint problem. BCP does not require any assumptions on the number of copy number states and computes the full posterior distributions rather than point estimates, reflecting the uncertainty in the locations of CNAs or copy number states. Furthermore, Erdman and Emerson made a 𝒪(*n*) MCMC implementation of the model, which allows for fast denoising, even for high-resolution data [5], and showed that BCP works well for segmentation of microarray data.

In this work, we introduce BayesCNA, a segmentation and copy number quantification method designed for low-quality liquid biopsy samples. BayesCNA utilizes information about posterior changepoint probabilities estimated using BCP, to extract likely postions of the CNAs in the genome. Changepoints in the genome that do not induce a significant change in the segment means are filtered away to avoid oversegmentation of the genome caused by the noisy nature of the data. We show that BayesCNA provides high-recall segmentation and accurate CNA detection using synthetic profiles and in silico sequencing results. Finally, we demonstrate the use of BayesCNA on experimental cell-line data where it achieves high concordance between detected CNAs in a baseline and low-quality replicates.

### 1.1. Outline of paper

The paper is organized as follows; In Section 2 we briefly introduce the Bayesian changepoint detection algorithm utilized in this study to infer changepoints, and how posterior changepoint probabilities are used to determine CNA positions in the genome. In Section 3, we evaluate the method using simulated and experimental data. The method is compared to the state-of-the-art methods QDNAseq [9] and ichorCNA [12] in terms of precision, recall and the F_1_-score, in low coverage and low purity scenarios. Finally, in Section 4 we discuss when BayesCNA should be used instead of conventionally used bioinformatical tools.

## 2. Methods

In this section, we introduce BayesCNA, a method for CNA detection and genomic segmentation that utilizes information about posterior changepoint probabilities estimated using BCP. Then, we discuss how CNA positions are determined using the posterior changepoint probabilities and how the signal is reconstructed based on this information. Finally, we discuss how synthetic data is generated and the experimental setup for the evaluation procedure.

### 2.1. Bayesian changepoint detection

Let 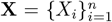 denote the data, where *X_i_* is the observed normalized copy number values (e.g. read depth information) in a genomic bin, and *n* the number of bins. The model assumes that there exists a partition of the genomic bins such that the mean is shared by all bins within each segment, corresponding to the copy number state of that segment. The observations are assumed to be independent 𝒩 (*μ_i_, σ*^2^), where *μ_i_* is segment specific but *σ*^2^ is shared throughout the genome. We assume that the noise in the samples is dominated by normal contamination, which is constant in the sample, making the assumption of equal variance reasonable. The prior of the mean *μ_ij_* of a segments spanning between positions *i* + 1 and *j* is assumed to be 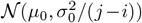, where *μ*_0_ and 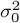 are hyperparameters. Note that the variance of the prior depends on the segment length, such that larger deviations from *μ*_0_ are expected in short blocks. This assumption is suitable for CNAs since small focal changes often show more copy number states and are also more affected by measurement noise. An exact implementation of Barry and Hartigan’s procedure is possible, but computationally expensive. Therefore, Erdman and Emerson [3] implemented an MCMC approximation and later reduced the computational complexity from 𝒪(*n*^3^) to 𝒪(*n*) [5]. For theoretical details and notes on prior specifications, we refer the reader to the original papers.

A partition is denoted as _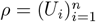_ where *U_i_* ∈ {0, 1} and *U_i_* = 1 is the indicator of a changepoint at position *i* + 1. Using MCMC, an approximate sample of *ρ* is generated in the following manner: initialize *U_i_* = 0 for all *i < n* and *U_n_* ≡ 1. A new partition is generated by iterating through the current partition, and at each position *i* assign *U_i_* = 1 with probability *p_i_* where

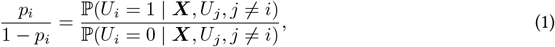

whose exact analytical form was derived by Barry and Hartigan [1] and simplified by Erdman and Emerson as incomplete beta integrals to ensure numerical stability for long sequences [3]. After iterating through the partition, the posterior means are updated conditionally on the current partition. Let **U** be a matrix whose rows *U* ^(*m*)^*, m* = 1*, …, M* consist of MCMC partitions with ^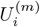^ being the indicator of bin *i* + 1 being a changepoint in partition *m* and *M* is the number of MCMC iterations after removing the burnin period. The posterior probability 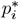 that a genomic bin *i* + 1 is a changepoint is estimated as

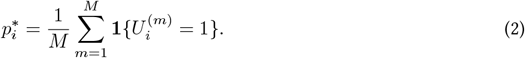

The method is implemented in the R package `bcp`by Erdman and Emerson.

### 2.2. Segmentation and postprocessing

In order to derive copy number profiles for downstream analysis from changepoint detection, segmentation is necessary, i.e. dividing the genome into segments of equal copy number states. We utilize the posterior changepoint probabilties to obtain a probable partition of the genome. Based on **U**, the most probable partition 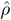can be estimated by computing the most frequent partition among the samples. However, we found that the frequency of each partition is low, due to the non-sharp boundaries in the data as a consequence of high noise levels, which reduced the accuracy of genomic segmentation. To circumvent this issue, we instead utilize information about the posterior changepoint probabilites for extracting likely positions of CNAs in the genome. We filter away noise in the posterior changepoint probabilities by setting

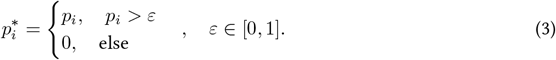

The parameter *ε* can either be fixed for all samples or chosen as, for example, 10% of the largest posterior changepoint probability, to account for differences between samples. When using such a cutoff, there is a risk of over-segmenting the genome, i.e., introducing too many changepoints and breaking up genomic segments with identical true copy number states. Especially in noisy situations, changepoints are found due to local trends in the data. Therefore, we undo changepoints that are not at least *η* standard deviations apart.

### 2.3. Copy number reconstruction through changepoint detection

In Figure 1 we outline BayesCNA and its required input and output. BayesCNA works for any binned genomic data, regardless of the preprocessing procedure applied. Briefly, we use the implementation by Erdman and Emerson [5] to generate a sequence of genome partitions and estimate the posterior changepoint probabilties and means. To gain insight into the uncertainty associated with the estimated mean, we also compute a 95% credible interval (CI) of the mean. Due to the noisy nature of the data, low changepoint probabilities are filtered away before running a peak detection algorithm to detect genomic bins of high posterior changepoint probability. After detecting high probability bins, the resulting copy number profile is taken as means or medians conditioned on the resulting partition. Changepoints introduced that do not result in a significant shift in mean (or median) are removed as these are expected to be a consequence of the noise. The method works as a preprocessing and denoising step, and the resulting signal can then be used in downstream analyses.

**Figure 1.**
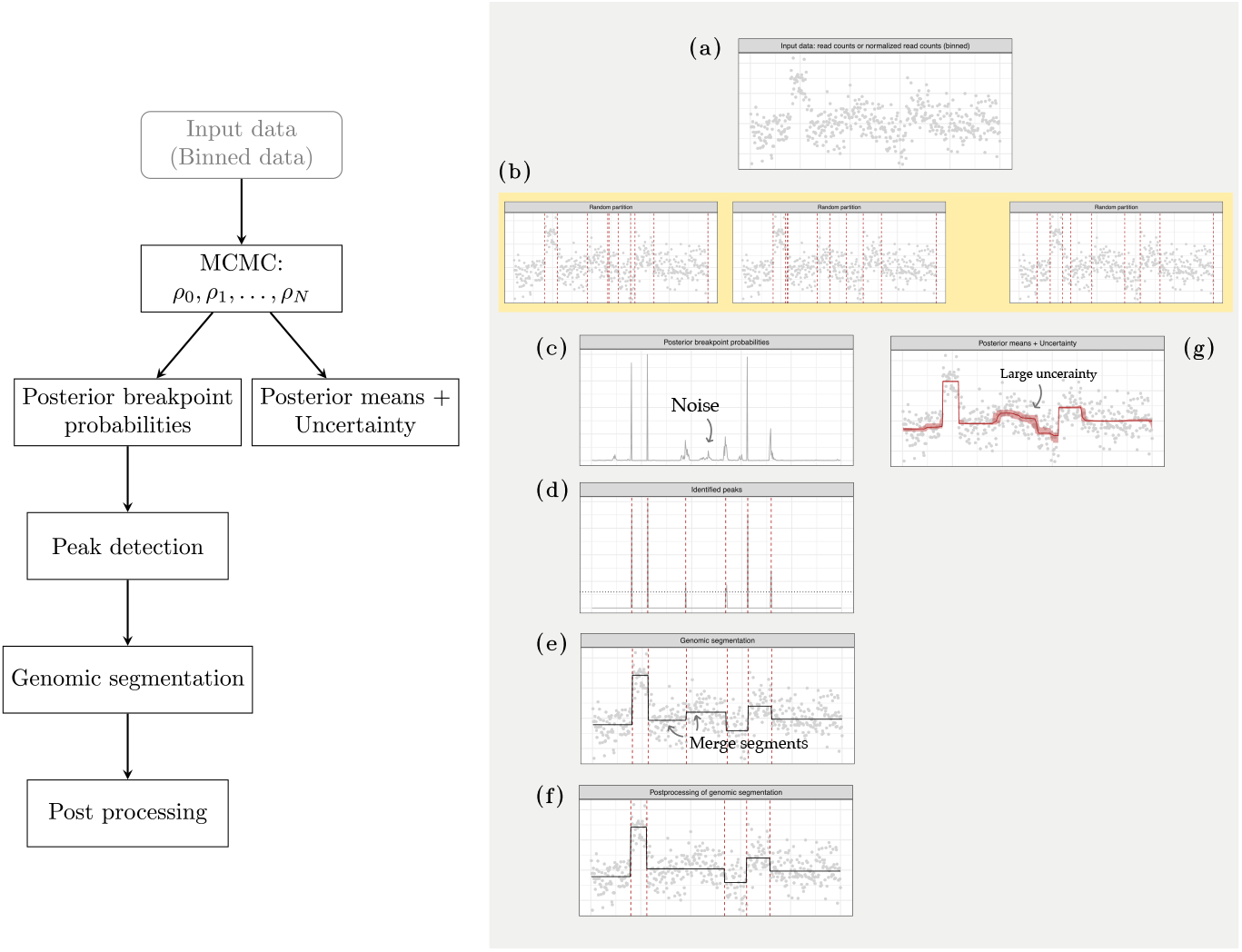
Outline of the estimation method BayesCNA. The right panel shows a flowchart of the individual steps and the right chart illustrates the corresponding steps. (a) The input data to BayesCNA is any genomic binned data. Here, the input data is simulated data of chromosome 1 with artificially introduced CNAs. (b) Examples of three random partitions generated in the MCMC step. The red dashed lines indicate the changepoints in the generated partition. (c) Posterior changepoint probabilities for each genomic bin. Low posterior changepoint probabilities are a consequence of noise in the data. (d) Posterior changepoint probabilities with identified changepoints marked out with the dashed red lines. In the peak detection algorithm, probabilities below a threshold *ε* are set to zero. (e) Genomic segmentation. Segment levels are computed as the conditional means based on the identified partition. (f) Post-processed genomic segmentation. Segment causing too small change in mean has been merged. (g) Posterior mean of the copy number profile and 95% credible interval. The intervals are computed using 2000 MCMC samples.

### 2.4. Simulation of synthethic and insilico datasets

We construct a synthethic dataset with artificial copy number profiles with *n* = 1000 genomic bins. For each sample, the segment length is drawn uniformly from *L* ∈ {30, 31*, …,* 50} and each segment is assigned a copy number state *C* ∈ {1, 2*, …,* 5} with probabilities 0.09, 0.5, 0.27, 0.09, 0.05 (rounded to two decimals). We control the sample purity *P* ∈ {0.05, 0.1*, …,* 0.5} by averaging the generated signal and a constant value of 2 (corresponding to contamination from healthy DNA), with weights *P* and 1 − *P* respectively. To reflect scenarios of different sequencing depths/coverages, we use four noise levels 0.5, 1, 2, 4, as described in [16]. Noise level 1 roughly corresponds to the normal amount of noise in an experimental sample, and the other noise levels represent a multiplicative factor times this baseline level. Furthermore, the noise is dependent on the copy number state, with larger copy numbers being associated with a larger noise level. Multiplicative noise is used to test the method in situations where the assumption of equal variance is violated.

An insilico dataset is generated by mixing simulated sequencing reads of tumor DNA and a healthy control. Raw sequencing data is created using CNVsim [19], a simulation software that can generate raw fastq-files using a predetermined set of alterations. We simulate artificial copy number profiles of chromosome 1 in a manner similar to the method described above. We ensure that at least 3 alterations are introduced and that the segment lengths of the alterations are in {30, 31*, …,* 50}. We avoid introducing any alterations in genomic bins {230, 231*, …,* 300} corresponding to the centromere (when using a 500kb bin size), as approximately this region will generally be removed by traditional bioinformatical tools. Furthermore, we ensure that the introduced alterations do not start or end within 10 genomic bins of this region and 20 bins within the start and end of the chromosome. The derived fastq-files are aligned to chromosome 1 of the human reference genome (version hg19, using bwa). To dilute the samples to the desired purity *P*, we mix simulated reads of tumor DNA (T) with reads of a healthy control (C), also provided by CNVsim. Let *N*_target_ be the desired number of reads for a fixed coverage level. The mixtures are constructed by first subsampling T and C to contain T_reads_ and C_reads_ respectively, such that T_reads_ + C_reads_ = *N*_target_, where

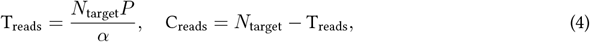

where *α* is the estimated purity of T, using liquidCNA [16]. Note that it is necessary to first estimate the purity of T, as tumor genomes are highly aneuploid often containing more DNA than a healthy genome, therefore our definition of purity (% of tumor cells contributing DNA) is not the same as the % of reads that have tumor origin. In the estimation of *α*, we use the fact that we have the ground truth available. In short, liquidCNA estimates purity by estimating the distance between peaks of the smoothed copy number distribution in a sample. The smoothing parameters of the density is chosen such that the number of peaks in the estimated density matches the true number of copy number states. Furthermore, the segment values in the estimation are taken as the median segment values, conditioning on the true partition. We discard the sample if the estimated purity is below 50%. The mixtures are then constructed by combining the reads using samtools merge. Note that the sample purity and coverage are approximate because of rounding errors in the subsampling step.

Finally, we construct a dataset of low-quality lpWGS data. We use lpWGS data derived from in vitro mixtures of two paired high-grade serious ovarian cancer (HGSOC) cell lines [13], as described in [16]. The samples are mixtures of a sensitive and resistant subclone and a healthy control of known proportions and have an average coverage of 1.3x. One technical replicate of the samples containing only tumour DNA (100% purity) are used to establish a baseline for the evaluation, as these samples naturally offer the most precise measurement of the true copy number states of the used cell line. Multiple samples are available to be used as baseline, corresponding to different sensistive/resistant mixtures. To construct the lowquality lpWGS data we subsample the data to contain approximately 10 million reads using samtools -s, corresponding to an average coverage of 0.15x. The samples are processed using QDNAseq in the same way as the baseline.

### 2.5. Processing of data

BCP is run using the `bcp`package, with 500 burnin iterations and *M* = 2000 MCMC iterations. We use the default *w*_0_ = 0.2 and set *p*_0_ = 0.01. This value of *p*_0_ was selected in an initial tuning stage as it works in both low and medium purity, but we suggest a smaller *p*_0_ if we know a priori that the sample is of higher coverage and/or purity, as this generally increases the precision, and a larger *p*_0_ if the sample is noisy to increase the recall. Furthermore, we use the threshold *ε* = 0.05 for filtering the posterior changepoint probabilities and *η* = 0.5 in the merging step, for the synthethic and insilico datasets. For the lpWGS dataset we use *ε* = 0.05 since diagnostic plots indicated that *ε* = 0.1 was too high.

We compare BayesCNA with the bioinformatical tools ichorCNA [12] and QDNAseq [9], which works on sequencing data on the insilico dataset. We run ichorCNA using the available snakemake workflow available from within ichorCNA, using bin size 500kb with the corresponding GC and mappability files, no matched normal panel, no clonality estimation, and standard parameters otherwise. For QDNAseq, genomic bins with mappability less than 75% are removed, in addition to the blacklisted regions. The normal sample is subsampled using samtools -s to contain the same number of reads as the mixture, and used to normalize the data.

### 2.6. Evaluation metrices

Let TP, FP, and FN be true positives, false positives, and false negatives, respectively. To evaluate the ability to detect copy number alterations, we compute the precision (positive predictive value; 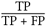) recall (sensitivity; 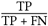) and the F_1_ -score defined as

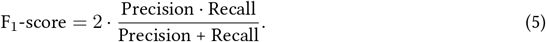

The F_1_-score combines precision and recall into a single metric and penalizes models with imbalances between precision and recall. We define a TP as a predicted changepoint that falls within ± 2 genomic bins within a true changepoint. If no changepoints are predicted (TP + FN = 0), we set the precision and the F_1_-score to zero to reflect the method’s inability to detect changepoints (otherwise, the precision and the F_1_-score is not defined). A predicted changepoint can only be assigned to a single true changepoint to prevent double counting of TPs, which would result in an inflated F_1_-score.

For the cell line lpWGS data, we instead study how much information is retrieved when reducing the coverage of the samples. We use the Normalized Mutual Information (NMI) metric:

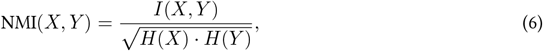

where *I*(*X, Y*) is the mutual information of two segmentations *X* and *Y*, and *H*(*X*) and *H*(*Y*) correspond to the entropy of *X* and *Y*, respectively. Intuitively, the mutual information *I*(*X, Y*) is a measure of the information that is shared between two variables *X* and *Y*. To obtain a fair measure regardless of the complexity of the variables, *I*(*X, Y*) is normalized to lie in the interval [0, 1]. If NMI = 0 there is no correlation between the samples, while NMI = 1 indicates perfect alignment. Here, *X* corresponds to a baseline and *Y* is the segmentation to compare with.

## 3. Results

Here, we evaluate BayesCNA on synthethic and experimental data. We compare BayesCNA to state-of-the-art bioinformatical tools ichorCNA [12] and QDNAseq [9]. ichorCNA uses a Hidden Markov Model (HMM) based method and a Bayesian inference framework, and QDNAseq uses the circular binary segmentation algorithm [2] for genomic segmentation.

### 3.1. Synthetic mixed populations

We begin by evaluating the performance of BayesCNA, on a dataset with known noise distribution (Figure 2). We generate synthetic data with varying levels of noise and purity values (Section 2.4). For each level of noise, we generate 50 samples per purity value {0.05, 0.1*, …,* 0.5}. Increasing levels of noise correspond to decreasing the sequencing depth/coverage. We observe that the predicitive ability decreases if either the purity or coverage is decreased, corresponding to an increase in the level of noise in the sample. We find a logarithmic like increase in the F_1_-score as a function of the purity. However, higher noise levels cause the growth of the F_1_-score to grow slower with increasing purity, because the signal is harder to distinguish from the noise in highly noisy scenarios. At noise level 0.5, the F_1_-score is close to perfect, except for purity 0.05 and some ouliers at higher purity levels. At noise level 1, we have an average F_1_-score greater than 0.75 with purities greater than 5% while a purity of 20% is required at noise level 2. Finally, we find that the method breaks down for the highest noise level and that sufficiently high purities are required to retain the predictive ability.

**Figure 2.**
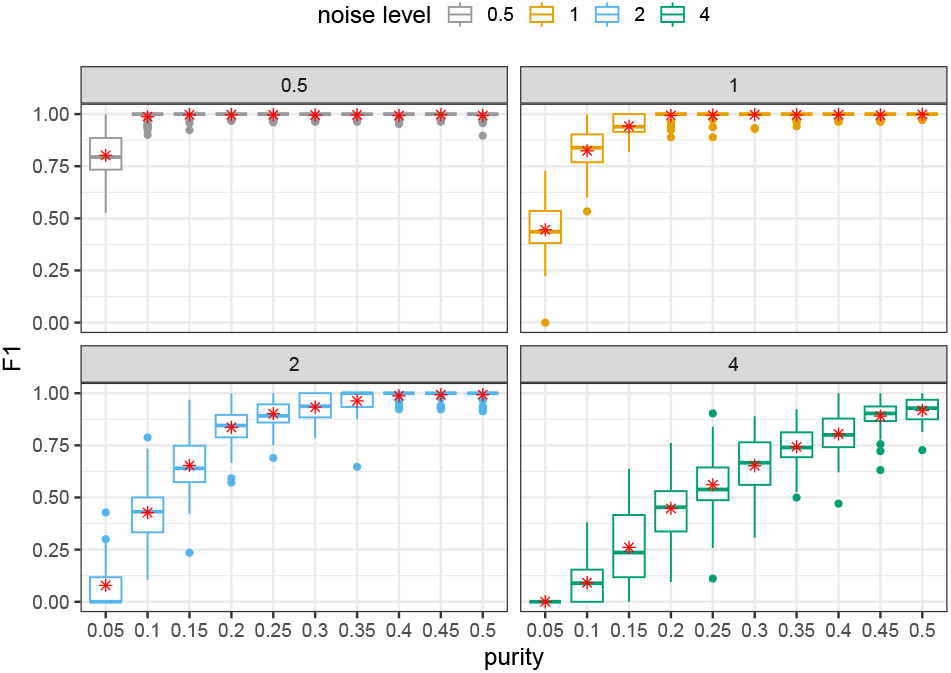
Predictive ability, measured as F_1_-score, of BayesCNA in synthetic mixed populations, with added multiplicative Gaussian noise. Increased noise levels represents decreased depth of coverage. The red stars in the boxplot indicates the mean of the distribution.

### 3.2. Insilico mixtures

Next, we compare BayesCNA to ichorCNA and QDNAseq in terms of precision, recall and the F_1_-score (Figures 3-4). We generate 20 samples of each combination of purity *P* ∈ {0.05, 0.1*, …,* 0.3} and coverage in {0.15, 0.2} (Section 3.1). To ensure a fair comparison, we run BayesCNA on the processed data produced by ichorCNA when comparing to ichorCNA, and on the data produced by QDNAseq when comparing to QDNAseq, as the normalization procedures differ between methods. In general, we find that the biggest advantage to using BayesCNA is in terms of recall, i.e. detecting more changepoints, especially for low-purity samples. The increase in recall is most evident when comparing the model with QDNAseq. Furthermore, the F_1_-score for BayesCNA is at least on par with the state-of-the-art tools. Figure 5(a) presents a case in which BayesCNA performs better than ichorCNA. Here, we find that BayesCNA is able to detect all changepoints, while ichorCNA is only able to detect the most prominent segment. ichorCNA also detects the start of the rightmost segments but fails to accurately identify the end of the segment. We suggest that BayesCNA should be used in low-quality cases where a more sensitive method is required to identify genomic alterations. In Figure 5(b), we instead present a case where ichorCNA performs better than BayesCNA. In this case, BayesCNA introduced too many alterations (lower precision) while ichorCNA correctly classified all CNAs. Hence, our method might be more sensitive to ‘intra-segment’ noise. However, noisy bins, or segments that are too short, can be removed in downstream analysis, while missed segmentation points cannot be recovered. We further note that the method was initially tuned to suit lowquality samples and that the selected parameters are suboptimal for high purity cases.

**Figure 3.**
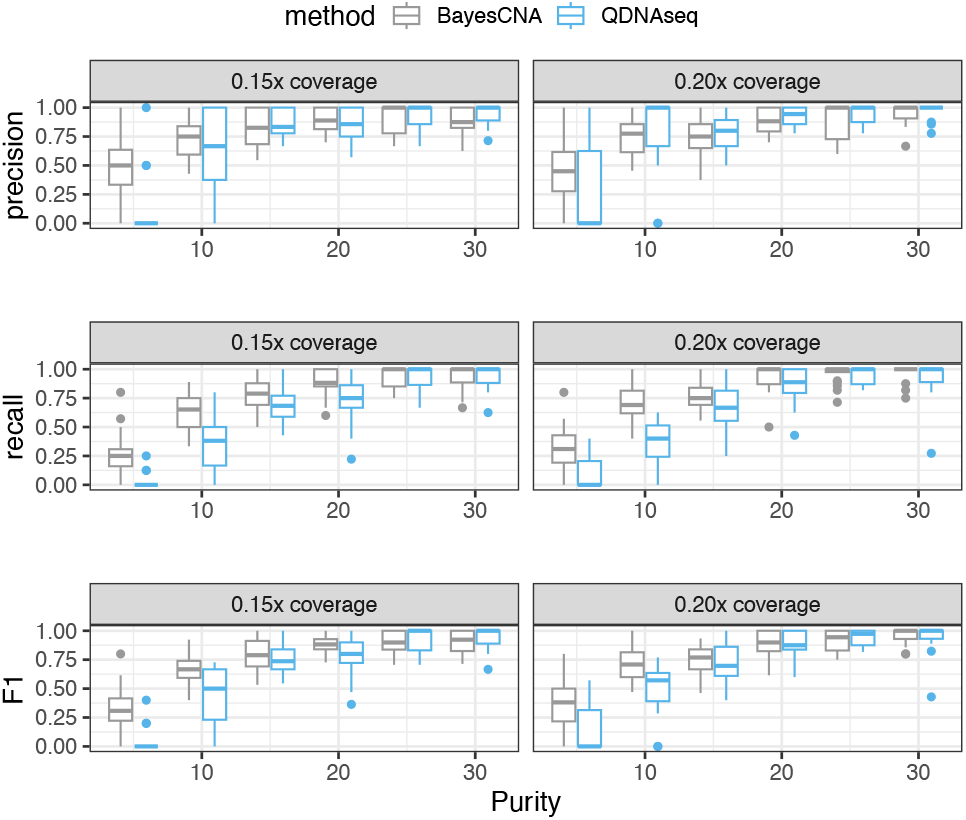
Comparison of changepoint detection ability between BayesCNA and ichorCNA, in terms of precision, recall, and the F_1_-score.

**Figure 4.**
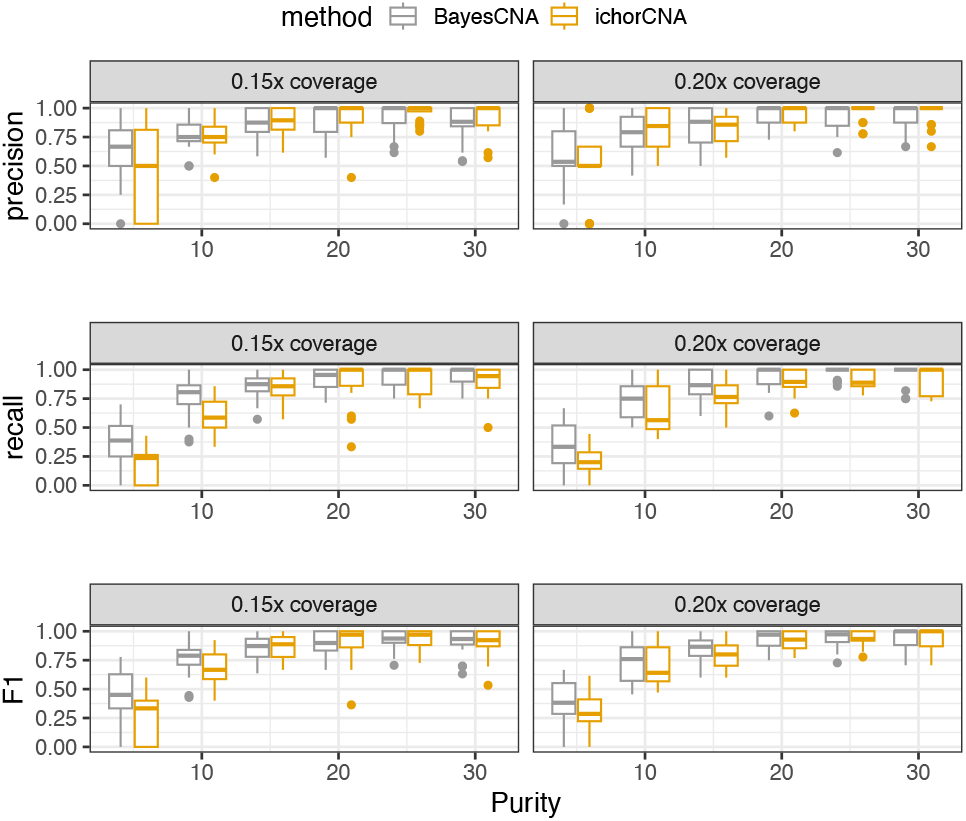
Comparison of changepoint detection ability between BayesCNA and QDNAseq, in terms of precision, recall, and the F_1_-score.

**Figure 5.**
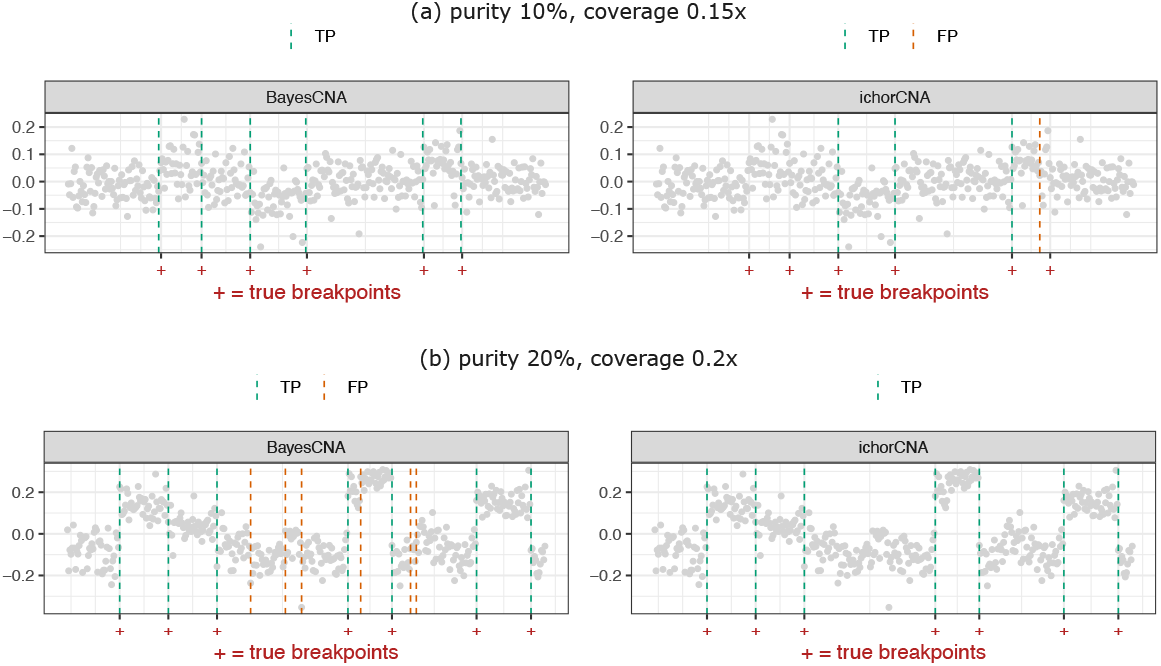
Illustrations of a case where BayesCNA performs (a) better, and (b) worse compared to state-ofthe-art bioinformatical tool ichorCNA.

### 3.3. Ovarian cancer cell line data

Finally, we evaluate the performance of BayesCNA on experimental data generated from ovarian cancer cell lines. As the true copy number states are unknown in this case, we instead rely on an established high-quality baseline sample, and measure the concordance, in terms of the NMI metric, between the segmentation of this baseline and low-coverage samples of the same DNA (Figure 6). As baselines for both BayesCNA and QDNAseq, we use the segmentation points detected in the low-noise and high-purity cell line sequencing data, as described in the Methods. We then evaluate how accurately the segmentation profiles are recovered from samples with lower tumor content and/or read count. We observe that for purities close to 20% (sample with approximately 81% purity not included), BayesCNA is on par with QDNAseq. In the most interesting region for this study, below 10%, we find that the NMI scores for BayesCNA are significantly higher compared to the NMI scores of QDNAseq, except for purity values close to 2%. However, we do not expect to extract any meaningful signal from samples with such low coverage and purity, since the inherent noise is on par with the signal. Furthermore, the method has not been tailored to such low purity values (these purity values were not included in the tuning and evaluation process). In conclusion, we find that BayesCNA retains more information when reducing the coverage of the samples, in the low purity region [3, 10], which are of interest in this study.

**Figure 6.**
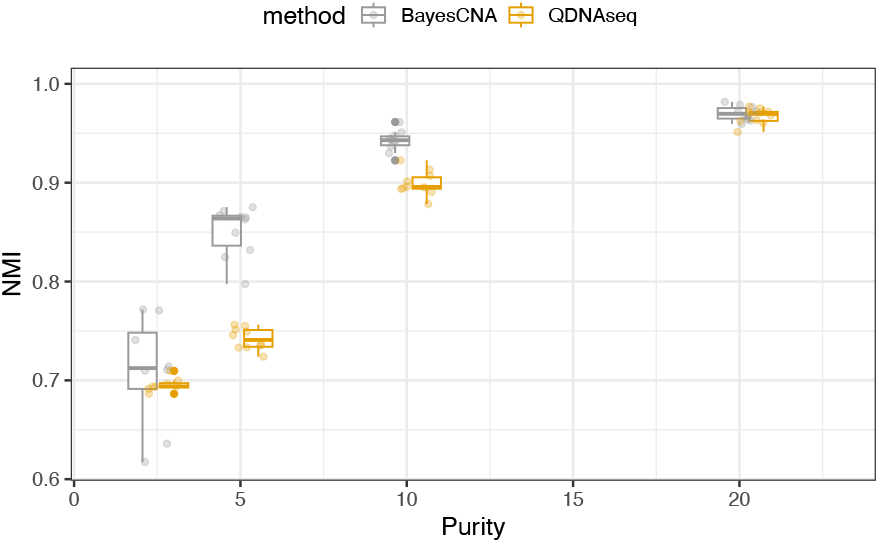
Comparison between QDNAseq and BayesCNA in terms of information retained when subsampling (increasing noise) in samples, in terms of the NMI metric.

## 4. Discussion

In this paper, we present BayesCNA, a computational method that detects CNAs and reconstructs the signal from low-coverage sequencing measurements from liquid biopsies. Our method uses Bayesian changepoint detection to estimate posterior changepoint probabilities for each position in the genome to infer likely start and ends of CNAs in the genome. A Bayesian approach is utilized because it naturally incorporates uncertainty, which is suitable for the noisy nature of liquid biopsy sequencing data. Using the detected partition of the genome, the signal is reconstructed by computing the segment means or medians conditionally on the partition. BayesCNA has been designed to increase recall (sensitivity) in low-coverage and low-purity samples where traditional methods struggle to detect any signal at all. BayesCNA works for any binned genomic data, regardless of the normalization procedure applied.

We validate the method’s ability to detect changepoints using both synthetic and insilico liquid biopsy sequencing measurements. We find that BayesCNA can detect changepoints with different types of noise distributions, even though it violates the assumption of the BCP model. When comparing BayesCNA to the traditional bioinformatic tools QDNAseq and ichorCNA, we find that BayesCNA offers higher recall, especially when compared to QDNAseq. We further validate the model in terms of the F_1_-score, where we find that the F_1_-score is at least on par with traditional methods. Finally, we demonstrate in experimental samples that we retain more information – quantified in terms of NMI – when increasing the noise levels, compared to QDNAseq. Since BayesCNA increases sensitivity, we suggest that BayesCNA can be a complement to traditional methods to increase the ability to detect CNAs where these methods tend to break down. Furthermore, it serves as a preprocessing step for methods that utilize copy number alteration profiles obtained from read depth information in their analyses. In particular, BayesCNA can be used upstream of methods relying large-scale CNAs or summary statistics across entire genomes in their analyzes, where larger alterations are often of most interest, with short segments potentially filtered away. On the other hand, BayesCNA might be less suitable for analyses working with high resolution and where the accuracy of short segments plays a large role in prediction.

We suggest that model parameters should be selected based on prior beliefs about the data or based on visual inspection of diagnostic plots. For example, if we know that the sample is of high purity, we should prefer a smaller *p*_0_ than used in this study and potentially filtering and merging of segments more aggressively. However, if the samples are of low quality, we should stick to less restrictive parameters to ensure that the signal is not lost in the post-processing. Furthermore, validation was only performed for alterations of length 30-50 genomic bins. We suggest that the parameters are tuned to suit the length distribution representing the sample at hand. We speculate that the method will generalize to longer segments, since this is generally a simpler problem, but shorter lengths require extra attention.

## 6 Author contributions statement

L.E. and E.L. conceived the study. L.E. designed the software, conducted the tests, and analyzed the results. L.E. and E.L. wrote and reviewed the manuscript.

## 7 Acknowledgments

We would like to thank Linnea Hallin for her work and input during the project leading up to this study.

## 9 Data and code availability

Aligned sequencing data from ovarian cancer cell lines is available from the European Nucleotide Archive (ENA: PRJEB42332). Raw CN values for these samples are available from https://github.com/elakatos/liquidCNA_data

All code implementing BayesCNA and analysing the results is available here: https://github.com/lottaer/BayesCNA

